# Genetic structure of *Rattus rattus* populations in an endemic plague focus in Madagascar: implications for rodent surveillance and management

**DOI:** 10.1101/2025.10.05.680582

**Authors:** Mamionah N. J. Parany, Anne Loiseau, Philippe Gauthier, Soanandrasana Rahelinirina, Gauthier Dobigny, Olivier Gorgé, Eric Valade, Minoarisoa Rajerison, Beza Ramasindrazana, Carine Brouat

## Abstract

**Background:** Plague remains a major public health concern in Madagascar. In the Central Highlands, where the disease is still endemic, the black rat (*Rattus rattus*) is the main reservoir of the causative agent *Yersinia pestis*. Understanding its population dynamics and structure is therefore crucial to inform control strategies, as dispersal may greatly limit the effectiveness of local interventions during outbreaks. In such a context, this study aims at investigating the genetic diversity and temporal structure of *R. rattus* populations at a fine geographical scale.

**Methodology/Principal findings:** Sampling was conducted in six villages of the Ankazobe district, both inside houses and outside villages. A total of 480 individuals, captured in March - May 2019 and 2020, were genotyped at 17 microsatellite loci. Our results show that genetic diversity levels were relatively homogeneous among villages and years. However, subpopulations living outside villages displayed significantly higher genetic diversity and lower genetic differentiation levels than those from inside houses, indicating larger effective population sizes outside villages in the cultivated habitats. These findings suggest more restricted movement among rat subpopulations from the houses, and greater connectivity among subpopulations living outside villages. However, overall genetic differentiation was rather low, suggesting extensive dispersal of rats at the scale of the district, facilitating rapid recolonization after local control efforts.

**Conclusion:** An integrated approach combining flea control within houses together with measures to reduce human-rodent contact would thus appear more appropriate than rodent control only to limit plague transmission.

## Introduction

Plague, though often perceived as a disease of the past, continues to pose serious public health challenges in certain regions of the world, particularly in Africa [1]. This zoonosis, caused by the bacterium *Yersinia pestis*, persists in natural foci. It affects a large variety of mammals, including rodents that act as reservoirs, while their fleas are vectors [2,3]. Because it circulates among many wild reservoir populations [4], plague cannot be eradicated [1]. Outbreaks of human plague usually occur within or at the vicinity of these natural foci, and their occurrence is partly related to the spatio-temporal dynamics of the reservoirs and vectors, and notably of commensal rodents and their fleas living in close contact with people [5–7]. Understanding these dynamics is therefore critical for improving surveillance and anticipating epidemic outbreaks on the one hand, and for implementing early effective control strategies on the other hand [8].

Madagascar is one of the major plague foci, with up to 80% of human plague cases reported worldwide annually [9]. The disease was introduced on the island over a century ago via maritime transports [10], and it rapidly moved to and became endemic in the central highlands, typically above 800 m high [11]. In rural areas of the highlands, the black rat (*Rattus rattus*) serves as the main reservoir of *Y. pestis* while two flea species, namely the cosmopolitan *Xenopsylla cheopis* and the endemic *Synopsyllus fonquerniei*, act as vectors [11]. Human cases are mostly reported during the hot and humid season, generally from September to April, a period corresponding to a low abundance of *R. rattus* [12] on the one hand, and to a peak of abundance of *S. fonquerniei* on the other hand [6]. During a plague outbreak, rodent and flea control is usually implemented at a local scale, i.e. inside and around houses of the hamlet where human cases have been reported, in order to reduce the populations of reservoirs and vectors. However, it has been argued that *R. rattus* is probably quite mobile and adaptable, often moving between habitats [13,14], which would make local control less effective.

Genetics can provide insights into spatio-temporal structure, population size and dispersal of reservoir populations, thus being very helpful in guiding rodent control strategies [15,16]. In Madagascar, previous studies based on microsatellite markers have shown that *R. rattus* population genetic structure may vary according to the relief in the central highlands, and may sometimes explain the distribution of human cases [17]. However, little is known about the genetic changes of rat populations through time at a fine geographical scale, something that would yet be relevant for plague prevention and control.

In this study, we investigated the population genetic structure of the main plague reservoir, *R. rattus*, within one district of the Central Highlands of Madagascar where plague is endemic and where human plague cases occur every year (Unpublished data, Central Laboratory for Plague, Institut Pasteur de Madagascar). We assessed rat population genetic diversity and estimated effective population size. We also investigated the temporal stability of genetic structuring and its association with habitat type (i.e., inside village houses *vs.* outside villages), with the aim of identifying the appropriate spatial scale for implementing rodent control measures.

## Materials and Methods

### Study areas and small mammals trapping

This study was carried out in Ankazobe district, covering a total area of 7,458 km^2^, and located in the central Highlands (mean altitude of 1,190 m) of Madagascar. Six localities were investigated, namely: Ambohitromby (18°25’S / 47°9’E), Ambolotarakely (18°1’S / 47°23’E), Antakavana (18°1’S / 47°23’E), Kiangara (17°54’S / 47°1’E), Talata-Angavo (18°12’S / 47°5’E) and Ankazobe I (18°12’S / 47°5’E) (Fig. 1). In these localities, villages are typically built on the top of hills, composed of 20 to 150 households and dwellings are mostly made of mud or brick with thatched roofs. Hills slopes are characterized by grassy areas while the lowlands (50 – 1,000 m away from the houses) are covered by rice paddies and food crops (vegetables, cassava or maize). The distance between localities varies from 10 – 80 km (Fig 1).

**Fig 1.**
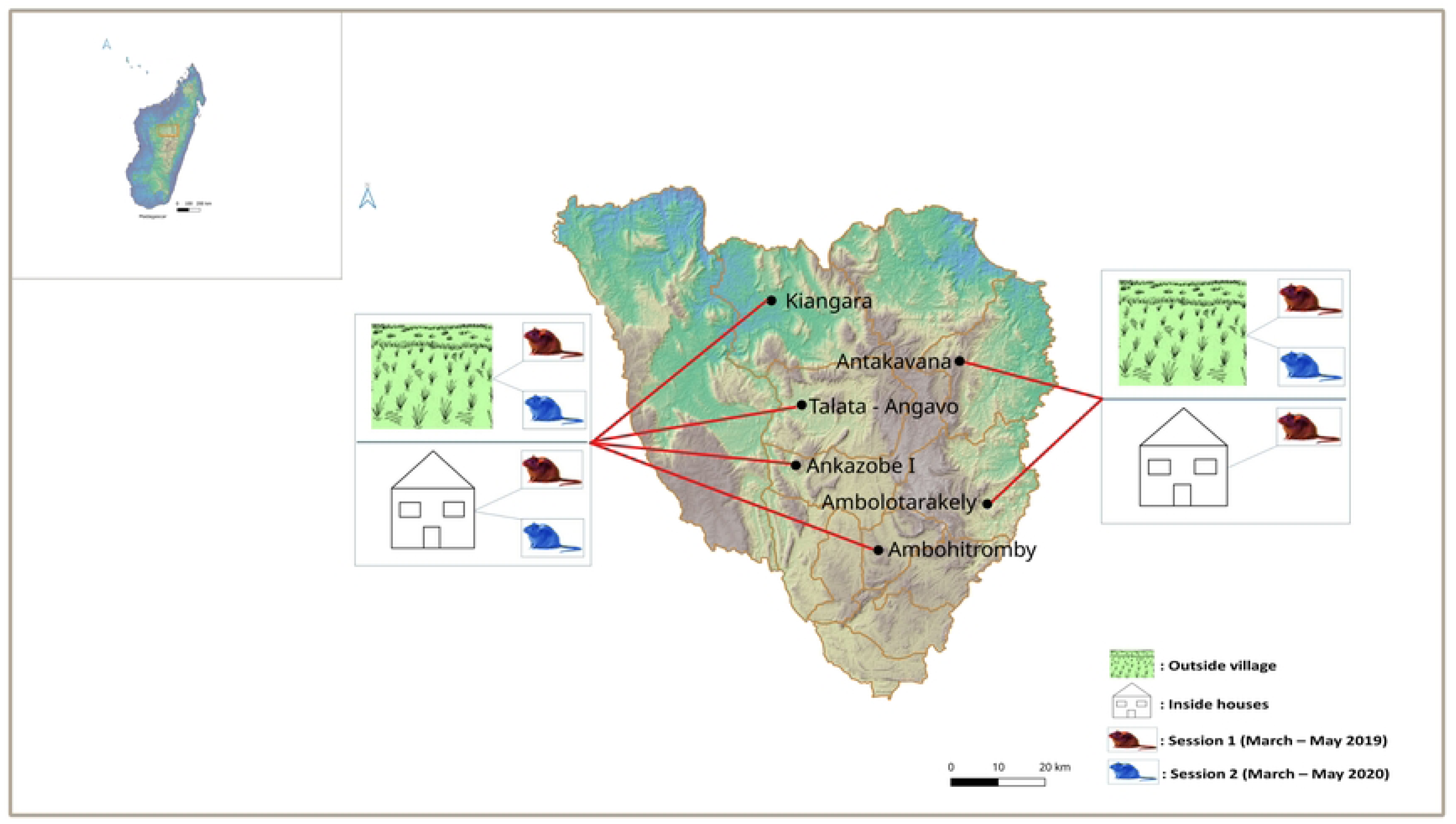
Sampling localities in Ankazobe district Trapping was performed in two habitat types (i.e., inside houses and outside villages) during two sessions. Topography data were obtained from NASA Shuttle Radar Topography Mission (SRTM) (2013). Shuttle Radar Topography Mission (SRTM) Global. Distributed by OpenTopography. https://doi.org/10.5069/G9445JDF.

Six small mammal Trapping Sessions (TS) were carried out in the six localities from December 2018 to June 2020. Trapping was systematically performed in two habitat types (i.e., inside houses and outside villages) in each locality (Fig 1). A detailed description of field protocols was already provided in Parany et al. (2025). For the present study, we focused on two capture sessions that were undertaken during the same period for two successive years: sampling 1 in March-May 2019, and sampling 2 in March-May 2020 (Fig 1). In each village, one BTS (Besançon Technique Service, 30L×10W×10H cm, BTS Company©, Besançon, France) and one Sherman (23L×7.5W×9H cm, H.B. Sherman Traps Inc©., Tallahassee, Florida, USA) traps were set inside 18 houses for three successive nights, thus reaching a total of 3,888 night-traps per village per session. Outside each village, 84 BTS traps were set for three successive nights (thus reaching 9, 072 night-traps per locality per session) following transect lines with 10-m intervals between traps along potential rodent pathways of reservoirs and/or near crop fields. All traps were baited every morning with a mixture of dried onions and dry fish, and checked twice a day. Traps that had captured rodents were replaced every morning.

### Data and sample collection

Each captured rodent was euthanized by cervical dislocation. Species, sex, external measurements (head-body length, tail length, ear length and hind foot length, in mm) and weight (in g) were recorded on each individual. Since terrestrial small mammals from the present experimental setting cannot suffer from potential taxonomic ambiguities, species identification could rely on phenotypic characteristics only (Soarimalala and Goodman 2011). Kidney of each individual was sampled then stored in 95° ethanol for subsequent molecular investigation purpose. Furthermore, blood was collected by cardiac puncture and centrifuged to obtain sera samples for serological analyses.

### Serological diagnosis of plague

Small mammal sera were tested for IgG antibodies specific to the *Y. pestis* F1 antigen using a well-established ELISA protocol [18]. Plague seroprevalence was then calculated as the percentage of seropositive individuals per locality, taking all individuals analyzed into account.

### Microsatellite genotyping

Molecular analyses were conducted on 11 to 40 *R. rattus* per habitat (i.e., inside houses or outside villages), locality and session. For each individual, total DNA was extracted from kidney using QIAamp DNA Mini Kit (Qiagen, Hilden, Germany) as recommended by the supplier. Microsatellite loci amplification was carried out in multiplex using a panel of 18 microsatellite markers, eight of which (namely, D10Rat20, D11Mgh5, D11Rat56, D16Rat81, D2Mgh14, D7Rat13 and D18Rat75) being originally developed for *R. norvegicus* (Jacob et al. 1995) while ten others (namely, Rr14, Rr17, Rr21, Rr22, Rr54, Rr67, Rr68, Rr93, Rr107 and Rr114) being *R. rattus*-specific [19]. Polymerase Chain Reactions (PCR) and genotyping were conducted according to previously described procedures [20,21]. Detection was conducted using an ABI 3130xl Genetic Analyzers (Applied Biosystems, Foster City, CA, USA) and genotypes were red, analysed and eye-cleaned under GeneMapper v.6 software.

### Data Analyses

Each group of samples from a given habitat × locality × session was considered as a distinct subpopulation in the subsequent analyses. Deviations from Hardy-Weinberg equilibrium (HWE) within subpopulations and genotypic linkage disequilibrium (LD) between pairs of loci were assessed using GENEPOP v.4 [22]. We corrected for multiple testing by the false discovery rate approach [23] implemented in the QVALUE package of R software v. 4.3.2 [24,25]. We used MICRO-CHECKER v.2.2.3 [26] to test whether heterozygote deficiencies could be accounted for by the existence of null alleles.

Genetic diversity of each subpopulation was estimated with FSTAT v.2.9.4 [27] by the allelic richness (*r*) calculated for a minimum of 10 individuals using the rarefaction procedure, *Nei’s* unbiased genetic diversity (*H_E_*, [28]) and *F_IS_*. Effective population size (*N_E_*) was estimated for each subpopulation with the LD methods using NE ESTIMATOR v.2.1 [29]. Alleles with a frequency ≥ 0.02 were used to minimise possible bias [30]. We compared *r*, *H_E_*, and *F_IS_*between habitats and between sessions using FSTAT (10,000 permutations), and *N_E_* using non parametric Kruskall-Wallis tests in R software v. 4.3.2 [25]. We investigated whether plague epizootics had left detectable traces on genetic diversity and effective population size by assessing in each locality the relationships between mean *H_E,_ r* and *F_IS_* on the one hand, and plague seroprevalence levels on the other hand using Spearman’s rank non parametric tests in R software v. 4.3.2 [25].

Genetic differentiation among each pair of subpopulations from each locality was summarized by calculating pairwise *F_ST_* estimates [31], with 95% confidence intervals (CIs) estimated by bootstrap resampling across loci using FSTAT v.2.9.4 (10,000 repetitions). *F_ST_*values were compared among habitats and sessions with FSTAT (10,000 permutations).

Genetic structure was further examined in several ways. First, we assessed whether locality may explain population genetic structure by performing an analysis of molecular variance (AMOVA; [32]) with ARLEQUIN v.2.000 (Schneider et al. 2000) using the locus-by-locus option. The variance components were tested using randomization (1,000 permutations). Second, we used G-based (log-likelihood ratio) randomization tests [33] to evaluate the effects of habitat (inside houses *vs*. outside villages) and sessions on genetic structure using FSTAT v.2.9.4 (10, 000 permutations of individuals between subpopulations for each analysis). Independent tests of pairwise genetic differentiation were combined using the generalized binomial procedure implemented in Multitest v.1.2 [34].

Under a model of isolation by distance (IBD), genetic distance between subpopulations is expected to increase with geographic distance. IBD was investigated by regressing pairwise estimates of *F_ST_* /(1 - *F_ST_*) against the logarithm of the Euclidean geographic distances between subpopulations [35], calculated using GPS coordinates. Mantel tests were performed to test the correlation between matrices of genetic differentiation and geographic distance in GENEPOP v.4 (10,000 permutations) for the whole dataset as well as considering each habitat separately.

Genetic structure was also examined using the Bayesian clustering approach implemented in STRUCTURE v.2.3.4 [36] in order to estimate the number of homogeneous genetic group (*K*) in our dataset. Analyses were performed with a model including admixture and correlated allele frequencies, for *K* ranging from 1 to10, using the LOCPRIOR option. Each run included 500,000 burn-in iterations following by 800,000 iterations. We performed 10 independent analyses for each *K* value. The number of genetic groups was inferred by the Delta *K* (ΔK) method applied to the log probabilities of data [37]. We also used implemented Discriminant Principal Component Analysis (DAPC; which can handle the absence of HW equilibrium) under R software v. 4.3.2 using *adegenet* and *devtools* packages in order to explore population genetic structure through an individual-based approach [38]. We determined the genetic groups (*K*) with the Bayesian Information Criterion (BIC), using the delta-BIC less than 6 [39,40] as a criteria. Both STRUCTURE and DAPC analyses were conducted on the whole dataset on the one hand, and on subpopulations sampled either inside houses or outside villages on the other hand.

### Ethics statements

This study is part of a murine surveillance program that aims at controlling plague reservoirs in Ankazobe district. Verbal authorization from the community’s authority and village heads, as well as verbal consent from the owners or tenants of investigated houses and fields were obtained after informing them about the project. Outside villages, traps were always set on the edge of cultivated fields, so as not to damage crops. Small mammals were treated in a human manner and manipulated by authorized and well-trained staff from the Plague Unit (Institut Pasteur de Madagascar) in accordance with guidelines of the American Society of Mammalogists (Sikes and the Animal Care and Use Committee of the American Society of Mammalogists 2016) and the directive 2010/63/EU of the European Parliament. In addition, all procedures used here for capture, autopsy and sampling of wild small mammals were validated by Institut Pasteur de Madagascar Animal Ethics Committee (agreement 001/2025/IPM/DS/CEA).

## Results

A total of 2,762 *R. rattus* were captured during the two years of sampling in the six localities of the Ankazobe District (see [12]). Among them, 480 individuals were genotyped for the purpose of the present study, corresponding to 11 to 40 individuals per habitat (inside houses vs. outside villages), locality and session, thus forming 22 subpopulations (Table 1). Rats captured inside houses during March-May 2020 in Antakavana and Ambolotarakely were excluded from the dataset due to low sample sizes (n = 6 and n = 7, respectively) (Fig 1).

**Table 1.** Genetic diversity estimates per subpopulations of *Rattus rattus* and global plague seroprevalence levels for each locality.

| Locality | Habitat | Session | N | $H_S$ | $r$ | $F_{IS}$ | $N_E$ | SP (%) |
| --- | --- | --- | --- | --- | --- | --- | --- | --- |
| **Ambohitromby (AMB)** | Outside villages | S1 | 40 | 0.72 | 6.2 | -0.007 | 370 [178 ; ∞] |  |
|  |  | S2 | 22 | 0.71 | 6 | -0.016 | 33 [26 ; 44] |  |
|  | Inside houses | S1 | 22 | 0.72 | 5.7 | 0.086 | 22 [18 ; 28] |  |
|  |  | S2 | 11 | 0.65 | 5 | 0.106 | 6 [3 ; 8] |  |
|  | **Overall** |  | **95** | **0.7** | **5.72** | **0.04** | **107.75 ± 131.125** | **0.37** |
| **Antakavana (ANT)** | Outside villages | S1 | 25 | 0.73 | 6.3 | 0.075 | 748 [156 ; ∞] |  |
|  |  | S2 | 21 | 0.73 | 6.3 | 0.076 | 147 [74 ; 1623] |  |
|  | Inside houses | S1 | 17 | 0.67 | 5.4 | 0.001 | 18 [14 ; 24] |  |
|  |  | S2 | NA | NA | NA | NA | NA |  |
|  | **Overall** |  | **63** | **0.71** | **6** | **0.05** | **304.333 ± 295.778** | **0.71** |
| **Ambolotarakely (ABK)** | Outside villages | S1 | 22 | 0.72 | 6.3 | 0.06 | 1301 [141 ; ∞] |  |
|  |  | S2 | 19 | 0.72 | 6.1 | 0.03 | 132 [64 ; ∞] |  |
|  | Inside houses | S1 | 23 | 0.72 | 6 | 0.129 | 13 [11 ; 15] |  |
|  |  | S2 | NA | NA | NA | NA | NA |  |
|  | **Overall** |  | **64** | **0.72** | **6.12** | **0.07** | **482 ± 546** | **0.21** |
| **Ankazobe I (ANK)** | Outside villages | S1 | 22 | 0.69 | 5.8 | 0.052 | 50[35 ; 84] |  |
|  |  | S2 | 21 | 0.7 | 5.8 | 0.106 | 42 [31 ; 64] |  |
|  | Inside houses | S1 | 22 | 0.68 | 5.2 | -0.043 | 10 [9 ; 12] |  |
|  |  | S2 | 18 | 0.69 | 5.9 | 0.093 | 35 [25 ; 53] |  |
|  | **Overall** |  | **83** | **0.69** | **5.68** | **0.05** | **34.25 ± 12.125** | **0.41** |
| **Kiangara (KIA)** | Outside villages | S1 | 21 | 0.74 | 6.3 | 0.037 | 113 [61 ; 542] |  |
|  |  | S2 | 22 | 0.72 | 6.2 | 0.037 | 49 [36 ; 73] |  |
|  | Inside houses | S1 | 17 | 0.7 | 5.2 | 0.073 | 11 [10 ; 13] |  |
|  |  | S2 | 22 | 0.69 | 5.5 | -0.017 | 15 [11 ; 19] |  |
|  | **Overall** |  | **82** | **0.71** | **5.79** | **0.03** | **47 ± 34** | **0.20** |
| **Talata-Angavo (TLA)** | Outside villages | S1 | 22 | 0.74 | 6.1 | 0.045 | 132 [69 ; 807] |  |
|  |  | S2 | 23 | 0.73 | 5.9 | 0.059 | 169 [80 ; ∞] |  |
|  | Inside houses | S1 | 22 | 0.72 | 6 | 0.066 | 27 [22 ; 34] |  |
|  |  | S2 | 22 | 0.69 | 5.5 | 0.163 | 20 [16 ; 25] |  |
|  | **Overall** |  | **89** | **0.72** | **5.89** | **0.08** | **87 ± 63.500** | **0.63** |

N: number of genotyped *R. rattus* individuals per subpopulations; $H_E$ : expected
heterozygosity; $r$ : allelic richness; $N_E$ : effective population size. SP: plague seroprevalence in
all small mammals sampled at the scale of the locality, NA: no estimate because of low sample size.

We excluded the locus D5Rat83 of the dataset because 49 individuals (10%) showed null genotypes following amplification problems. All other loci were found at HW equilibrium after correction for multiple testing, except D11Rat56 that displayed significant heterozygote deficiencies in several subpopulations, probably due to null alleles. The estimated mean frequency of null alleles at this locus was 20%, which may slightly impact genetic diversity and differentiation estimators [41]. We verified for most analyses that results were similar with or without this locus, and presented in the following the results obtained using this locus. After correction for multiple testing, LD was not significant in any of the tests performed. So, the 17 loci were considered to be independent.

Genetic diversity estimates were relatively homogeneous among subpopulations. Allelic richness (*r*) ranged from 5 to 6.3 alleles (mean 5.9 ± 0.1), and expected heterozygosity (*H_E_*) from 0.65 to 0.74 (mean 0.71 ± 0.01) (Table 1). Mean *F_IS_* values ranged from 0.03 to 0.08 (Table 1). Furthermore, genetic diversity estimates were significantly different among habitats, with subpopulations outside villages being slightly more diverse than subpopulations inside houses (*r*: *p* =0.001; *H_E_*: *p* = 0.01; no significant difference for *F_IS_: p* = 0.27). They were not different among sessions (*r*: *p* = 0.75; *H_E_*: *p* = 0.29 and *F_IS_: p* = 0.57).

Effective population size *N_E_* values ranged from 6 to 35 for inside houses subpopulations, and 33 to 1,301 for subpopulations outside villages, with an overall mean of 157 **±** 178 (Fig 2). There were significant differences between subpopulations (Kruskal Wallis test: χ^2^ =15.135, df = 1, *p* < 0.001), *N_E_* being higher for subpopulations sampled outside villages than for those living inside houses (Fig 2).

**Fig 2.**
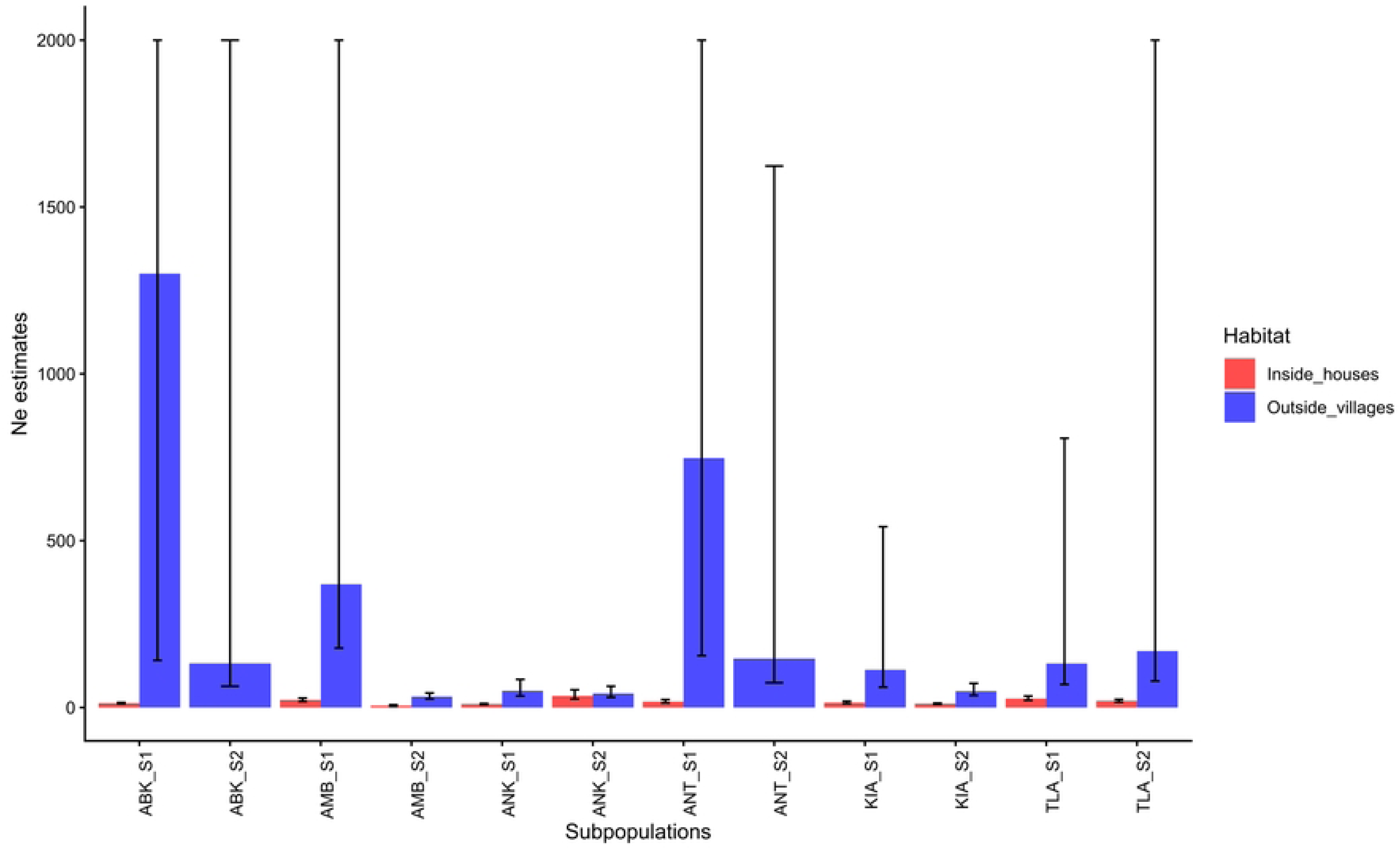
Mean *N_E_* and its standard deviation, calculated using the LD method, in each subpopulations of *Rattus rattus* sampled inside houses and outside villages of the six localities during two trapping sessions (S1 and S2). ABK: Ambolotarakely; AMB: Ambohitromby; ANK: Ankazobe I; ANT: Antakavana; KIA: Kiangara; TLA: Talata-Angavo.

The overall plague seroprevalence across all localities was low (0.4%), ranging from 0.2% to 0.71% (Table 1). No significant correlation was observed at the locality level between plague seroprevalence (calculated from our dataset of 2,762 small mammals trapped over the entire 2-year long survey) and mean *H_E_* (Spearman’s ρ = −0.25*, p* = 0.65), *r* (ρ = 0.08, *p* = 0.91) and *F_IS_* (ρ = 0.49*, p* = 0.32).

The mean *F_ST_* value computed across all subpopulations was 0.032 (95% CI = [0.028; 0.037]). Pairwise *F_ST_* estimates ranged from 0.0002 to 0.38 among all subpopulations. They were significantly higher among subpopulations sampled inside houses than among subpopulations outside villages (*p* = 0.007), but similar among subpopulations sampled in one session or in the other. The AMOVA analysis showed significant genetic differentiation between localities (*Va* = 2.08%; *p* < 0.001) although most of the genetic variation was observed within individuals (*Vc* = 90.46%; *p* < 0.001). There was also significant genetic differentiation among habitats within localities (unweighted mean *F_ST_* = 0.095; generalized binomial test *p* <0.001). Genetic differentiation among sessions within localities was lower but also significant (unweighted mean *F_ST_*= 0.003; generalized binomial test *p* < 0.001).

Genetic IBD was not significant among subpopulations (*p* < 0.0001; slope *b* = −0.001, 95% CI = [−0.005, 0.003]). However, there was a tendency towards a positive IBD when considering only subpopulations sampled inside houses (*p* = 0.10; slope *b* = 0.058, 95% CI = [0.02, 0.098]).

STRUCTURE was used to investigate whether rat populations where spatially structured in Ankazobe district. The highest *ΔK* value (26.87) was rather low, indicating no substantial substructure at the scale of the district, but weak differentiation between northern and southern subpopulations (Fig 3A). This differentiation between northern (Kiangara, Antakavana, Ambolotarakely and Talata-Angavo) and southern (Ambohitromby and Ankazobe I) subpopulations was more pronounced when only subpopulations inside houses were considered (Fig 4A). Alternatively, no substructure was observed when only subpopulations outside villages were analyzed. The DAPC analyses confirmed the results obtained with STRUCTURE, indicating no substructure at the scale of the whole dataset (Fig 3B), and a slight South-North substructure when only individuals sampled inside houses were taken into account (Fig 4B).

**Fig 3.**
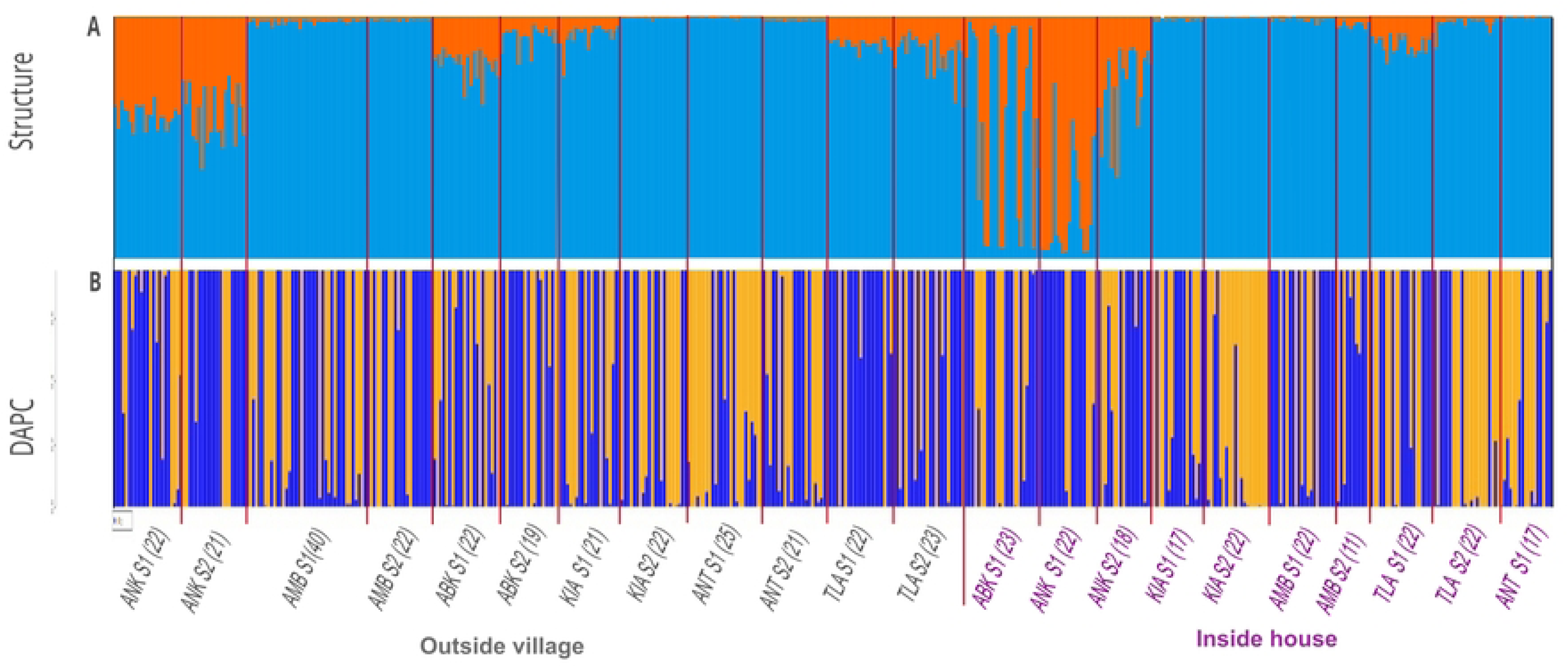
Genetic structure of *R. rattus* for all subpopulations (outside villages and inside house) of the Ankazobe district analyzed with STRUCTURE (A) and DAPC (B) with K=2. In the barplots, each individual is represented by a vertical bar with the colors showing the proportion of the individual genotype derived from respective genetic clusters. ANK: Ankazobe I; AMB: Ambohitromby; ABK: Ambolotarakely; KIA: Kiangara; ANT: Antakavana; TLA: Talata-Angavo.

**Fig 4.**
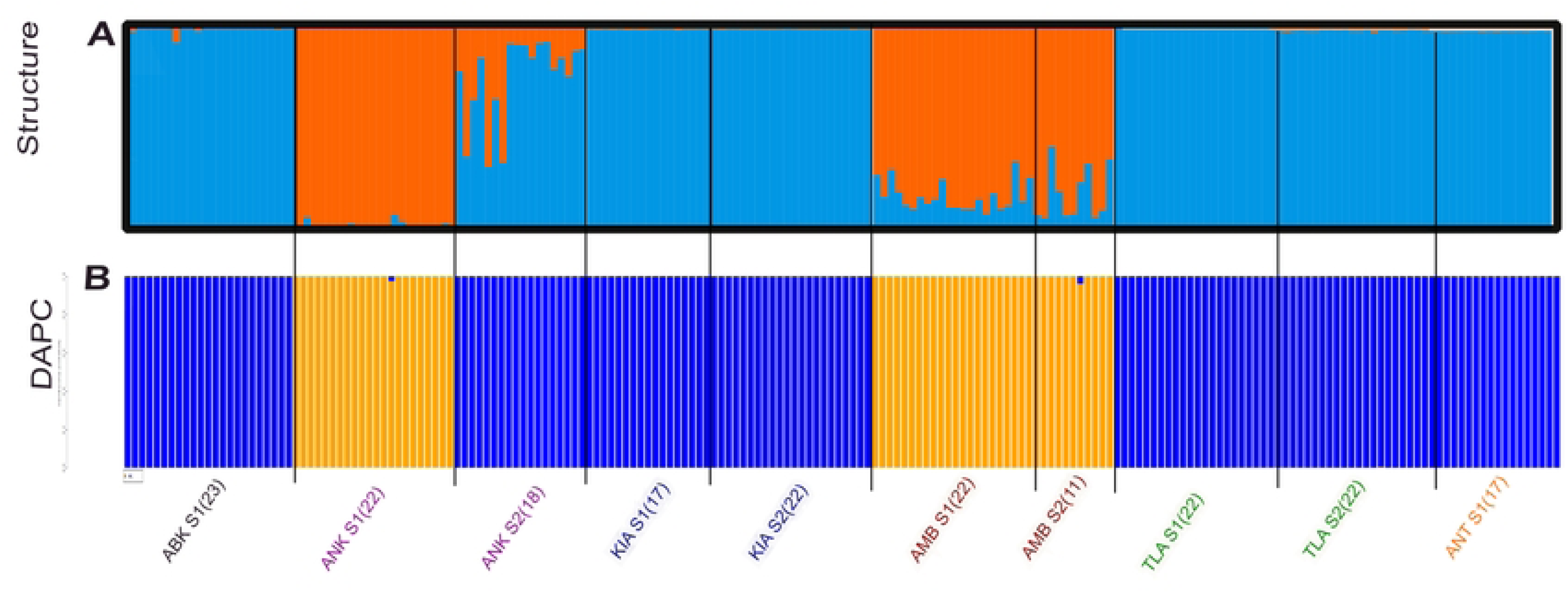
Genetic structure of *R. rattus* for all subpopulations sampled inside houses of the Ankazobe district, analyzed using STRUCTURE (A) and DAPC (B). See legend of Fig. 3 for more information.

## Discussion

In Madagascar, in addition to the urban foci in the surroundings of Mahajanga seaport (West coast) [42,43], two main long-lasting plague foci have been described: the first one in the northern central area of the country, and the second one in the Central Highlands of Madagascar [11]. Ankazobe district is part of the latter one where plague remains endemic with suspected human plague cases being detected every year [44,45]. Previous studies have shown that *R. rattus* is the primary reservoir of *Y. pestis* and triggers human outbreaks in this district [12,45]. They were consistent with those of previous studies demonstrating the implication of *R. rattus* as the reservoir of plague in the Central Highlands of Madagascar [11,46].

During our 2 year-long survey, rather low seroprevalence was observed in small mammals, suggesting a limited circulation of *Y. pestis* in the six localities studied [12]. However, human plague cases were regularly recorded during the same period (2019-2020) in other localities of the district. As such, from 2019 to March 2020, a total of 19 cases were reported, of which seven were confirmed at the Central Laboratory for Plague of the Ministry of Health (Unpublished data, Central Laboratory for Plague, Institut Pasteur de Madagascar). Low seroprevalence levels could also be related to the fact that a substantial proportion of *R. rattus* from the Central Highlands was shown to be resistant to plague, some of them (up to 30%) remaining seronegative after experimental exposure [47,48]. Previous studies conducted in other plague-endemic regions of Madagascar also reported a low number of seropositive rats, even despite frequent human outbreaks, which was interpreted as consequence of their plague resistance [17].

Genetic diversity estimates were relatively homogeneous among subpopulations of rats in the six localities and between both sessions. They were relatively high (mean *r* = 5.9, mean *H_E_* = 0.71) when compared to those observed in other African settings, for instance in Cotonou, Benin (*r* = 4.4, *H_E_* =0.58: [49]), in Niamey, Niger (*r* = 3.4, *H_E_* =0.5: [50]) or in Dakar, Senegal (*r* = 3.9, *H_E_* =0.67: [51]). Nevertheless, they were similar to those observed in other localities of the Central Highlands of Madagascar (mean *r* between 4.8 and 5.5, and mean *H_E_* between 0.68 and 0.72 in four areas studied in [17]. Such differences between Malagasy and other African populations of black rats may suggest contrasted population sizes, which would be larger in Madagascar than in the other continental areas where the black rat is restricted to the commensal environment (Cotonou, Dakar and Niamey) and sometimes distributed in spatial patches (e.g., Niamey). They may also reflect differences in bio-invasion history, the black rat having been introduced much earlier in Madagascar (probably from the 11^th^ century: [52]) than in West Africa (probably during the European colonization era: [51,53]).

Genetic diversity was slightly higher for subpopulations outside villages compared to those from inside houses. This difference was consistent with that observed for effective population sizes (*N_E_*). The larger sizes of subpopulations outside villages may reflect the much larger and continuous habitat, while habitat inside villages is of limited size (i.e., houses and their immediate surroundings). The temporal stability of genetic diversity estimates over one year (which may represent up to 2-3 generations of rats) and their lack of correlation with plague seroprevalence (as evaluated at the locality scale) suggests that plague epizootics, if any, did not leave any significant effect on population sizes. Alternatively, this impact may be too limited in intensity (relative to population size) or in time length, to impact genetic diversity [54].

Genetic differentiation among *R. rattus* subpopulations was globally low, indicating dispersal of individuals at the scale of the Ankazobe district, and/or large population sizes. This is consistent with previous results obtained in four other areas of the Malagasy highlands, reporting globally low genetic differentiation levels in black rat populations at similar spatial scales, which however increased with topographic relief [17]. Despite this low level of differentiation, our results suggest that subpopulations of inside houses and outside village are differentiated within one given locality (unweighted mean *F_ST_*= 0.095), indicating limited effective dispersal among habitats. Limited movements of black rat individuals between both habitats would be consistent with the observation of different ectoparasites on rats sampled in fields (mainly parasitized by *Synopsyllus fonquerniei*) compared to those sampled within houses (infested by *Xenopsylla cheopis*) [55]. While rats reproduce all year round in houses probably following permanent food resources inside houses, they reproduce seasonally outside villages [12,56]. Seasonal reproduction could imply longer-distance movements of individuals living outside villages than those living inside houses, thus potentially explaining lower genetic differentiation among subpopulations sampled outside villages. On the contrary, individuals living inside houses would have less need to move in order to find mates or food, thus leading to slightly lower mobility. This hypothesis is supported by the study of Rahelinirina et al. (2010), which used Rhodamine B to mark rats and track their movements in rural Madagascar: they showed that marked rats inside houses were primarily recaptured near their initial marking points [57].

The north-south pattern found by STRUCTURE and DAPC analyses in subpopulations sampled inside houses may reflect isolation by distance, which may be only marginally significant due to the limited number of sampled localities. Alternatively, it could be driven by differences in population spatio-temporal dynamics: southern localities appear to be more directly linked to major urban centers and could thus be more likely colonised by rats coming from localities outside the Ankazobe district. Also, southern localities reported higher numbers of human plague cases compared to others localities of the district between 2014 and 2018 (Unpublished data, Central Laboratory for Plague at IPM). Such high numbers of human cases may have been preceded by plague epizootics and subsequent recolonization of local populations by black rat individuals from different origins.

Finally, population genetics results may be useful to define management/eradication units and to guide control strategies [58]. In Madagascar, local rodent control is usually recommended from March to August, before the onset of the plague season, in order to reduce the number of potentially breeding individuals and to prevent subsequent increase in rat abundance. In Ankazobe, however, our results demonstrate the existence of widespread and highly connected populations of *R. rattus* at the scale of the district, which would seriously challenge the effectiveness and lasting effects of such local rodent control campaigns. In this context, an integrated approach combining rodent as well as flea control within houses, together with environment-based management towards reduction of human-rodent contact may appear as a much more appropriate strategy to limit plague circulation among rats on the one hand, and plague transmission from rat to human on the other hand.

## Conclusion

This study revealed that *R. rattus* populations in Ankazobe district exhibit high genetic diversity, low differentiation and inter-annual stability. Significant differentiation between subpopulations sampled inside houses *vs.* outside villages reflect different spatio-temporal patterns in the two types of habitats. Altogether, our results questioned the efficiency of local rodent control when used alone to fight against plague. Rather, they advocate for more integrated strategies taking into account rodents, flea vectors and rodent-flea-human contacts.

## Acknowledgment

We would thank the team of the Plague Unit of the Institut Pasteur de Madagascar for their help in field and laboratory works associated with this study. We sincerely thank the Centre de Biologie pour la Gestion des Populations (CBGP), Montpellier France for its valuable support in carrying out the laboratory work and for its guidance in the genetic analyses. Microsatellite genotyping was done at the molecular biology platform of the CBGP and through the GenSeq technical facilities of MEEB (CNRS and University of Montpellier) hosted by ISEM (CNRS, University of Montpellier and IRD). We are also grateful to the technical teams in the different study sites during the fieldworks. We thank also Institut Pasteur de Madagascar for the financial support to MNJP (Institutional PhD grant, Bourse Girard).

## Author contributions

Conceptualization and design were conducted by MNJP MR CB BR EV OG; methodology by MNJP CB AL SR PG; formal analysis by MNJP CB GD; the original draft was written by MNJP CB, with all authors providing comments to produce the final version of the manuscript.

